# Evolutionary changes in the number of dissociable amino acids on spike proteins and nucleoproteins of SARS-CoV-2 variants

**DOI:** 10.1101/2023.03.12.532219

**Authors:** Anže Božič, Rudolf Podgornik

## Abstract

The spike protein of SARS-CoV-2 is responsible for target recognition, cellular entry, and endosomal escape of the virus. At the same time, it is the part of the virus which exhibits the greatest sequence variation across the many variants which have emerged during its evolution. Recent studies have indicated that with progressive lineage emergence, the positive charge on the spike protein has been increasing, with certain positively charged amino acids improving the binding of the spike protein to cell receptors. We have performed a detailed analysis of dissociable amino acids of more than 1400 different SARS-CoV-2 lineages which confirms these observations while suggesting that this progression has reached a plateau with omicron and its subvariants and that the positive charge is not increasing further. Analysis of the nucleocapsid protein shows no similar increase of positive charge with novel variants, which further indicates that positive charge of the spike protein is being evolutionarily selected for. Furthermore, comparison with the spike proteins of known coronaviruses shows that already the wild-type SARS-CoV-2 spike protein carries an unusually large amount of positively charged amino acids when compared to most other betacoronaviruses. Our study sheds a light on the evolutionary changes in the number of dissociable amino acids on the spike protein of SARS-CoV-2, complementing existing studies and providing a stepping stone towards a better understanding of the relationship between the spike protein charge and viral infectivity and transmissibility.

## INTRODUCTION

The COVID-19 pandemic caused by the SARS-CoV-2 virus gave rise to an unprecedented global effort to collect and share the information of the virus’ ongoing evolution [1, 2] with its emerging stream of new variants with varying degrees of disease severity, transmissibility, and other traits [3, 4]. Consequently, this large amount of data enables the study of how the virus has been changing since it was first detected in humans and the discovery of those mutations which made it adapt further, with a particular focus on the designated variants of interest (VOI) and variants of concern (VOC) [5–8]. The spike (S) protein of the virus—responsible for target recognition, cellular entry, and endosomal escape [9]—in particular exhibits the greatest sequence variation, not only across different SARS-CoV-2 variants but also in coronaviruses in general [10, 11]. As an example, the highly transmissible omicron variant has 15 mutations solely in its receptor-binding domain (RBD), the part of the S protein which binds to angiotensin-converting enzyme 2 (ACE2) receptor. The high variability of the S protein across different variants has also led to recent efforts in studying whether VOCs could be identified from the sequence of the S protein alone, which would make lineage assignment relatively easier compared to the use of complete genome sequences [12].

Numerous experimental, computational, and bioinformatics studies have analyzed the influence of the observed mutations in the S protein on the viral function. Some of the mutations have been linked to an increased transmissibility of the virus, while others have been linked to changes in its binding to ACE2 [6, 17–19]. A particularly interesting observation, made as the COVID-19 pandemic progressed, concerns the tendency of the mutations in the S protein of different VOCs to increase the number of positively charged amino acid (AA) residues in it [20] (illustrated on a few examples in Fig. 1). This observation is further supported by a more recent analysis of the charge on the S protein of major SARS-CoV-2 lineages which identified a striking change in the S protein charge with the evolution of the virus [21]. Molecular dynamics and large-scale ab initio studies also found that the increase in the (partial) charge of S protein RBD of some VOCs increases the binding to ACE2 and other cell surface receptors [14, 22–25] and that the total charge of the RBD might be a simple predictor for the RBD–ACE2 binding affinity based on the data obtained for main VOCs [26]. That charge might influence viral infectivity has also been shown in avian influenza viruses, where hemagglutinin proteins from low and high pathogenic strains exhibit clear difference in their surface charge [27]. These observations go hand in hand with a large body of studies showing the electrostatic interactions to be of enormous importance both in biological systems in general [28, 29] as well as in viruses in particular [30, 31].

**Figure 1.**
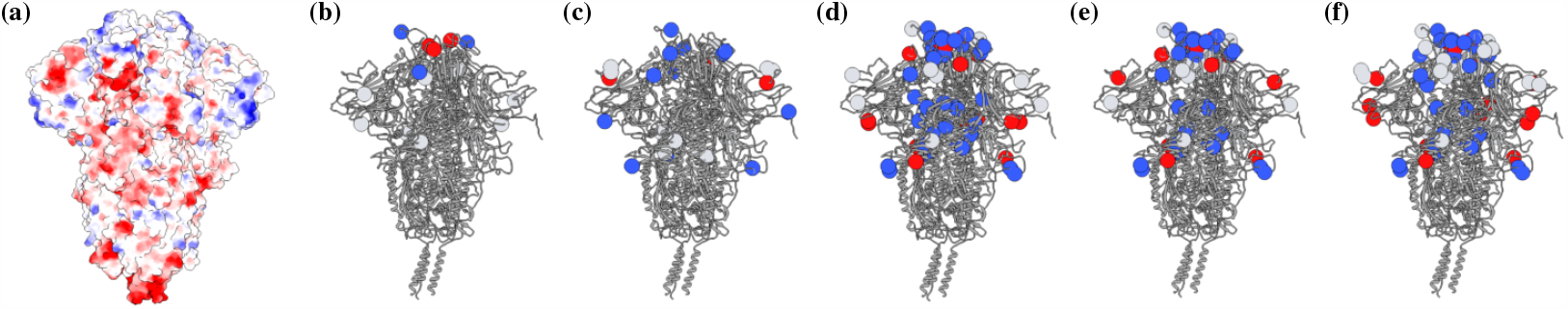
**(a)** Coulombic surface colouring of the WT SARS-CoV-2 S protein (PDB: 7FB0), showing the positions of positively and negatively charged AAs (blue and red, respectively) on the S protein surface. The actual charge distribution on the S protein is a more complex question which depends, among other things, on the bathing solvent conditions, such as electrolyte concentration and pH, and local values of AA dissociation constants [13, 14]. **(b)**–**(f)** Mutations of dissociable AAs on the S protein of selected SARS-CoV-2 VOCs: beta (b), delta (c), omicron BA.1 (d), omicron BA.4 (e), and omicron XBB (f). Only those mutations that lead to either a gain or a loss of a dissociable AA are shown [15]: gain of a positively charged AA (blue), gain of a negatively charged AA (red), and loss of either positively or negatively charged AA (light gray). Mutations which replace one dissociable AA with another of the same charge type are not represented. Note that these images show only the positions of the mutations of dissociable AAs and do not depict the changes in charge on the S protein. The complete PDB structure of the S protein in panels (b)–(f) was obtained from the work of Kim *et al*. [14]. All the images were drawn with UCSF Chimera [16].

To study the effect of S protein charge on, for instance, its binding affinity, requires significant computational effort and more often than not involves a certain degree of approximation [13, 14, 26], unless one resorts to quantum-level calculations [32]. On the other hand, the amount of dissociable AAs provides a good estimate for the total charge of the S protein and importantly enables a broad study of different SARS-CoV-2 lineages [20, 21]. In this work, we use the large amount of available data on different SARS-CoV-2 lineages that have emerged over the past three years of the pandemic to examine in detail how the number of dissociable AAs on the S protein—as a proxy for its total charge—has changed with increasing lineage divergence and evolution of the virus. We observe several different clusters corresponding to the emergence of different variants. We indeed observe a general tendency towards an increase in the total number of positively charged AAs on the S protein, a tendency which, however, seems to have recently plateaued. For comparison, we perform the same analysis on the SARS-CoV-2 nucleoprotein (N protein) to show that the evolutionary preference for positive charge is specific to the S protein. We additionally examine the available data on S and N proteins in known coronaviruses to frame our results in a wider context. In this way, our work complements the existing studies on the importance of S protein charge for its interaction with the environment, at the same time adding several novel observations regarding the preference for particular dissociable AAs in different variants and the saturation of their number with omicron VOC and its many subvariants.

## RESULTS

### Dissociable AAs on S and N proteins show different amount of change with lineage divergence

In general, there are six AAs which can (de)protonate and thus acquire charge: three of them can be positively charged (arginine (ARG), lysine (LYS), and histidine (HIS)), whereas three of them can be negatively charged (aspartic (ASP) and glutamic (GLU) acid and tyrosine (TYR))—see also Methods. Figure 2 shows how the average number of different dissociable AAs on the S and N proteins of 1421 SARS-CoV-2 variants changes with increasing average lineage divergence from the wild-type (WT) genome. Since the length of the S protein is approximately three times the length of the N protein (*≈* 1270 AA compared to *≈* 415 AA, respectively), it is not surprising that the number of dissociable AAs on the S protein is in general much larger than the number of dissociable AAs on the N protein. One can nonetheless observe very different trends in their numbers as lineage divergence increases. For instance, the number of positively charged LYS on the S protein tends to steadily increase with lineage divergence, with a peak number of around 67 with the omicron subvariants BA.1, BA.3, and the recombinant XD (Fig. 2a). However, this number slightly decreases afterwards for the more divergent subvariants BA.4, BA.5, and the recombinant XBB—see also Fig. 3 and Table I. The number of HIS also increases with increasing divergence, albeit it does so only at relatively highly divergent lineages. The changes in the number of negatively charged AAs are less prominent, with perhaps the exception of TYR, whose number is slightly increased with the more divergent omicron subvariants.

**Figure 2.**
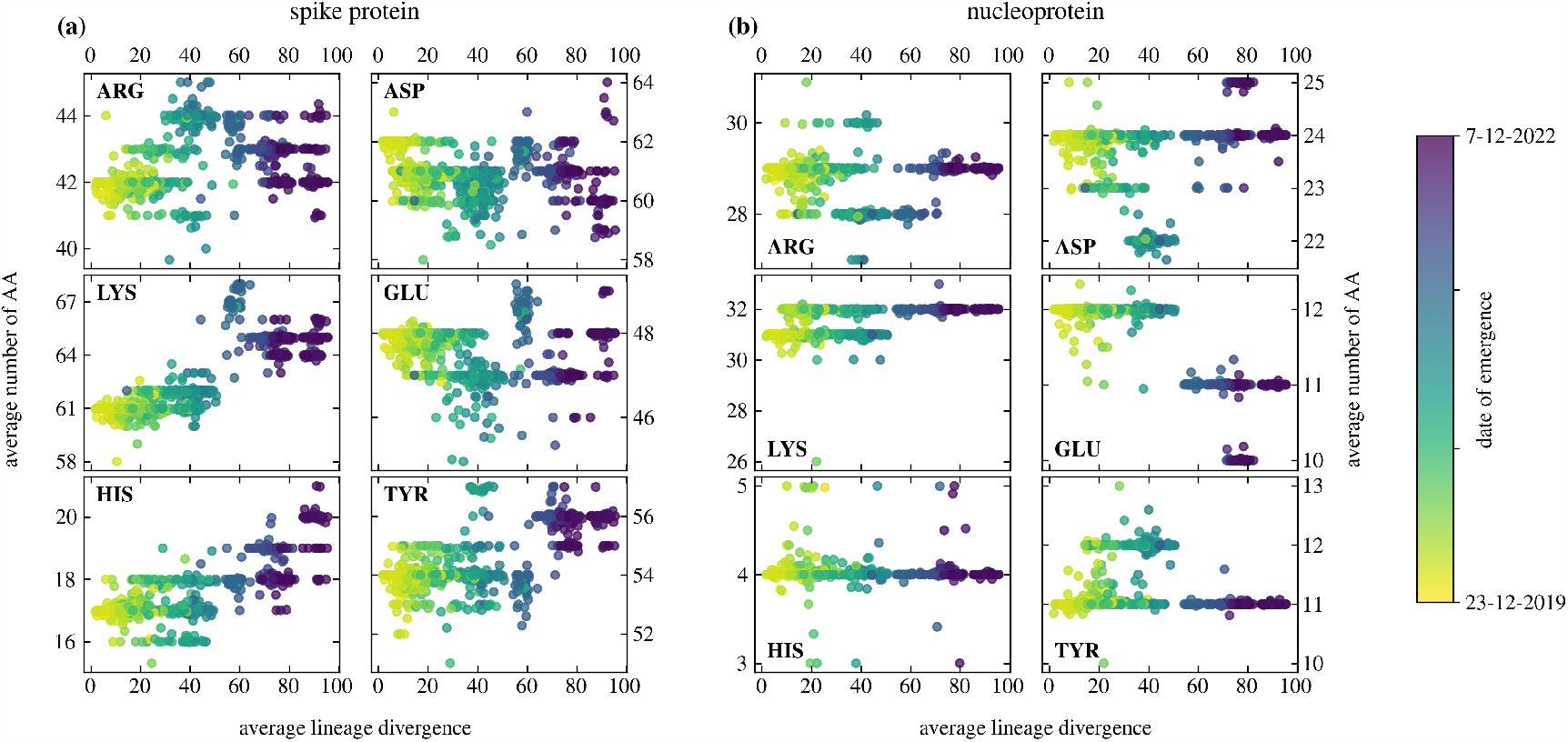
Average change in the number of dissociable AAs on **(a)** the S protein and **(b)** the N protein of 1421 different SARS-CoV-2 lineages as a function of average lineage divergence. Left column of each panel shows positively charged AAs (ARG, LYS, HIS) and the right column of each panel shows negatively charged AAs (ASP, GLU, TYR). Datapoint colours correspond to the earliest known isolation date of the lineage as indicated by the colourbar.

**Figure 3.**
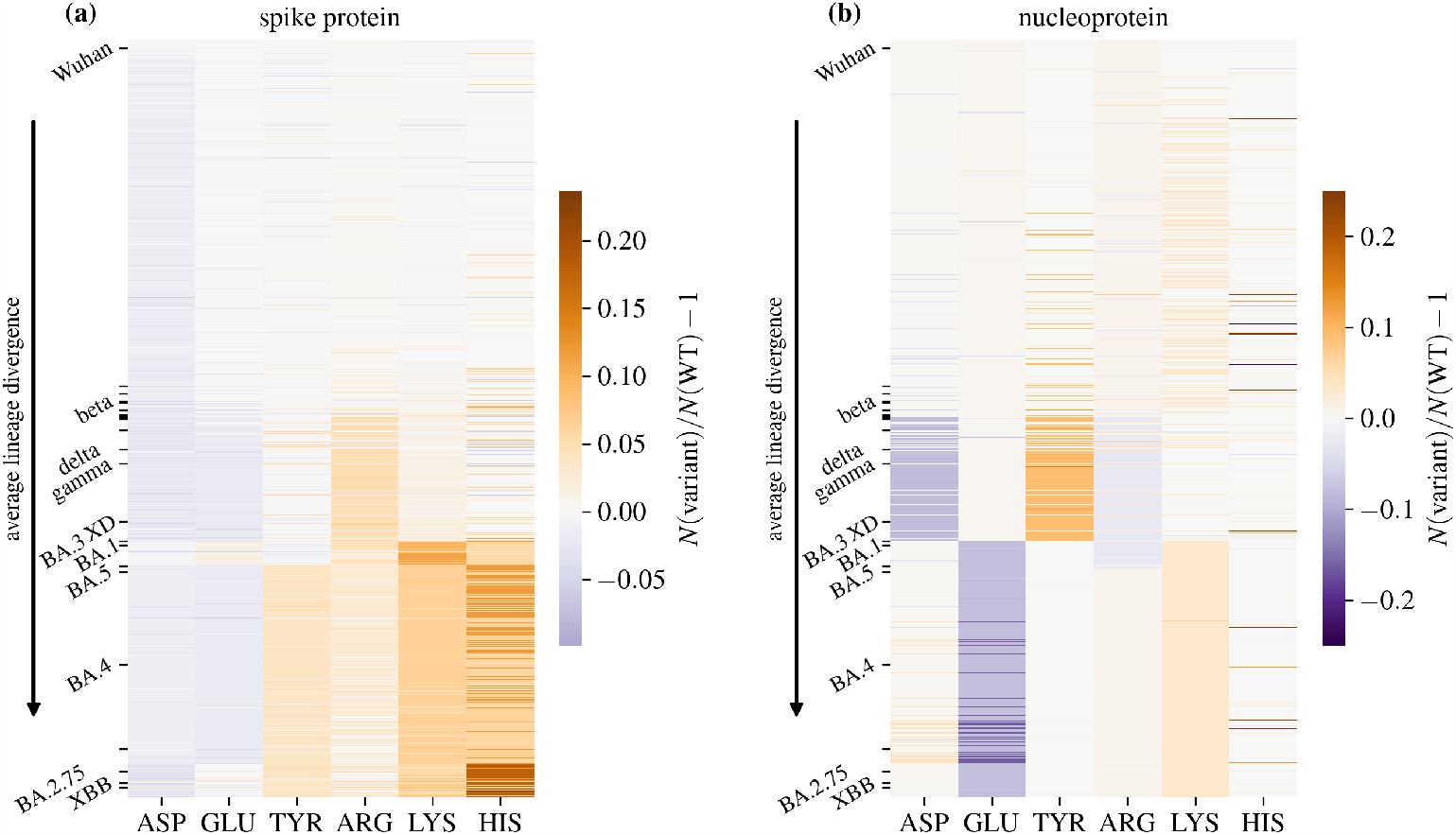
Heatmap of the relative change of the average number of dissociable AAs on **(a)** the S protein and **(b)** the N protein of 1421 SARS-CoV-2 lineages compared to the WT. Lineages are sorted in the order of increasing divergence. Ticks and labels on the *y* axes mark select VOIs and VOCs along the divergence progression (cf. Table I).

**Table I.** Average lineage divergence of selected SARS-CoV-2 VOCs and VOIs.

| Variant | Pango lin. | Avg. divergence |
| --- | --- | --- |
| Wuhan | B | 5.02 |
| iota | B.1.526 | 25.2 |
| zeta | P.2 | 25.8 |
| beta | B.1.351 | 27.7 |
| epsilon | B.1.429 | 28.1 |
| eta | B.1.525 | 30.5 |
| kappa | B.1.617.1 | 32.5 |
| theta | P.3 | 33.0 |
| lambda | C.37 | 33.8 |
| mu | B.1.621 | 36.9 |
| delta | B.1.617.2 | 38.8 |
| gamma | P.1 | 39.7 |
| omicron XD | XD | 44.4 |
| omicron BA.3 | BA.3 | 54.9 |
| omicron BA.1 | BA.1 | 56.1 |
| omicron XE | XE | 63.7 |
| omicron BA.5 | BA.5 | 68.3 |
| omicron BA.4 | BA.4 | 73.5 |
| omicron XAY | XAY | 79.0 |
| omicron BA.2.75 | BA.2.75 | 88.7 |
| omicron XBB | XBB | 91.1 |
| omicron BJ.1 | BJ.1 | 92.0 |

In contrast to the S protein, the number of dissociable AAs on the N protein does not show any significant increases or decreases with lineage divergence (Fig. 2b). Here, the more interesting observation is that the number of certain AAs, such as TYR and LYS, occupied two distinct values in the early variants. As the lineages began to diverge more, only one of the values is selected for (a lower number of TYR and a higher number of LYS, respectively). Nonetheless, the number of dissociable AAs on the N protein shows far fewer changes with divergent SARS-CoV-2 lineages compared to their number in the S protein.

### Different positively charged AA types are preferred with lineage divergence

As Fig. 2 shows, different dissociable AAs show different patterns of change with the evolution of SARS-CoV-2 and the emergence of new lineages. In order to see whether any patterns can be observed between different lineages with respect to their preference for a particular AA type, Fig. 3 shows a heatmap of the relative changes in the number of dissociable AAs on the S and N proteins from different SARS-CoV-2 lineages compared to the WT, with select VOCs and VOIs marked along the divergence progression. Looking at the S protein first (Fig. 3a), we can observe that the first positively charged AA residue to show an increase is ARG, which reaches a slight peak in a cluster of variants which covers delta and gamma variants (cf. also Table I). The number of ARG, however, decreases again for more divergent variants, and in its place, the number of LYS is significantly increased, in a cluster covering omicron subvariants BA.1, BA.3, and XD. While the number of LYS also remains high for later, more divergent subvariants, its number nonetheless drops in comparison to this cluster. Interestingly, the most divergent variants, including the omicron subvariants B.2.75, BJ.1, and the recombinant XBB, show an increased number of HIS, which is only fractionally charged at physiological pH. The numbers of negatively charged AAs on the S protein change less drastically, but have a tendency towards a slight decrease in the more divergent lineages, with the notable exception of TYR, the number of which is increased in most omicron subvariants.

While the overall number of dissociable AAs on the N protein changes less drastically (Fig. 3b), some trends can nonetheless be observed. Variants such as delta and gamma show an increase in the number of TYR and a decrease in the number of ASP. On the other hand, these changes are absent in the omicron subvariants, where the most notable observation is the decrease in the number of GLU. We again observe that as lineage divergence increases, the number of positively charged AAs on the N protein shows a much lesser degree of change compared to their counterparts on the S protein. Figures 3 and 2 together make it clear that the evolutionary preference for the increase in the amount of positively charged AAs is particular to the S protein of SARS-CoV-2, and no similar effect can be observed for the number of dissociable AAs on the N protein.

### Number of positively charged AAs on the S protein has plateaued with omicron subvariants

As already mentioned, the total number of dissociable AAs can serve as a proxy for the amount of charge on the S and N proteins. Figure 4 separates the contributions of the positively (ARG, LYS, HIS) and negatively (ASP, GLU, TYR) charged AAs and shows how their total number changes with the increasing divergence of SARS-CoV-2 lineages from the WT genome. Similarly to what has been observed previously [20, 21], the number of positively charged AAs on the S protein increases with lineage divergence, while the number of negatively charged AAs remains rather steady or even decreases slightly for highly divergent lineages (Fig. 4a). This indeed indirectly implies that the overall charge on the S protein has been becoming more positive with increasing lineage divergence. However, we can also observe that the total number of positively charged AAs appears to have reached a plateau with the omicron variant, with only small changes in the amount of positively charged AAs observed between its different subvariants.

**Figure 4.**
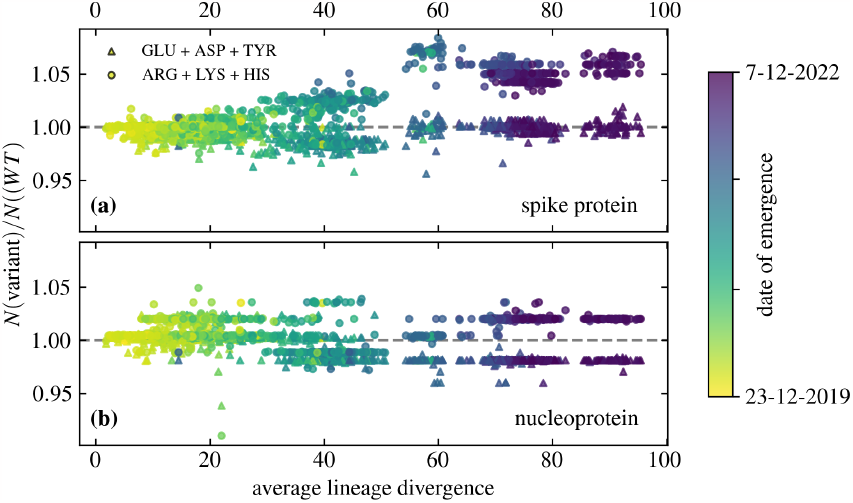
Change in the average total number of AAs on **(a)** the S protein and **(b)** the N protein of 1421 different SARS-CoV-2 lineages compared to the WT, shown as a function of the average lineage divergence. AAs are grouped by their ionizability: ASP, GLU, and TYR (negative) on the one hand and ARG, LYS, and HIS (positive) on the other.

The changes in the total number of positively and negatively charged AAs on the N protein as lineage divergence increases are, on the other hand, again far smaller compared to those of the S protein (Fig. 4b). However, we can also observe that the most divergent (and recently emerged) lineages seem to clearly prefer a version of the N protein with slightly fewer negatively charged AAs and slightly more positively charged AAs compared to the WT, albeit with a significantly smaller variation in their number compared to the less divergent lineages. This, combined with the observations of the number of charged AAs on the S protein (Fig. 4a), implies that the number of dissociable AAs on both the S and N proteins has reached an “equilibrium” where any further significant changes appear to be less likely.

### SARS-CoV-2 has more positively charged AAs than other known (beta)coronaviruses

Compared to the abundance of data on SARS-CoV-2, there is much less information available regarding the evolution of the S proteins of other coronaviruses. Nonetheless, we can compare the number of dissociable AAs on the S protein of most currently known coronaviruses based on their reference sequences. Figure 5 thus shows the comparison of the number of positively and negatively charged AAs on the S proteins of the reference strains of 46 different coronaviruses (see Methods), together with 22 VOCs and VOIs of SARS-CoV-2 (Table I). In general, the S proteins of coronaviruses tend to have a larger number of GLU and ASP compared to the number of ARG and LYS (Fig. 5a), as well as a large amount of TYR (Fig. 5b), which indicates that the overall charge of the S protein is negative. There are a few exceptions which have approximately the same number of GLU and ASP compared to ARG and LYS, including human coronavirus NL63, suncus murinus coronavirus X74, two avian coronaviruses (IBV and 9203), and two bat coronaviruses (rhinolophus bat coronavirus HKU2 and rousettus bat coronavirus HKU9). Compared to most other betacoronaviruses—and, in fact, most other coronaviruses in general—both WT SARS-CoV-2 and its variants have a larger number of ARG and LYS on their S proteins. One notable exception is SARS-CoV, which has a similar amount of positively and negatively charged AAs as WT SARS-CoV-2. In general, Fig. 5 shows that the S protein of WT SARS-CoV-2 already has fewer negatively charged AAs and more positively charged AAs than other betacoronaviruses and that with lineage divergence, it is the number of positively charged AAs that has increased even further.

**Figure 5.**
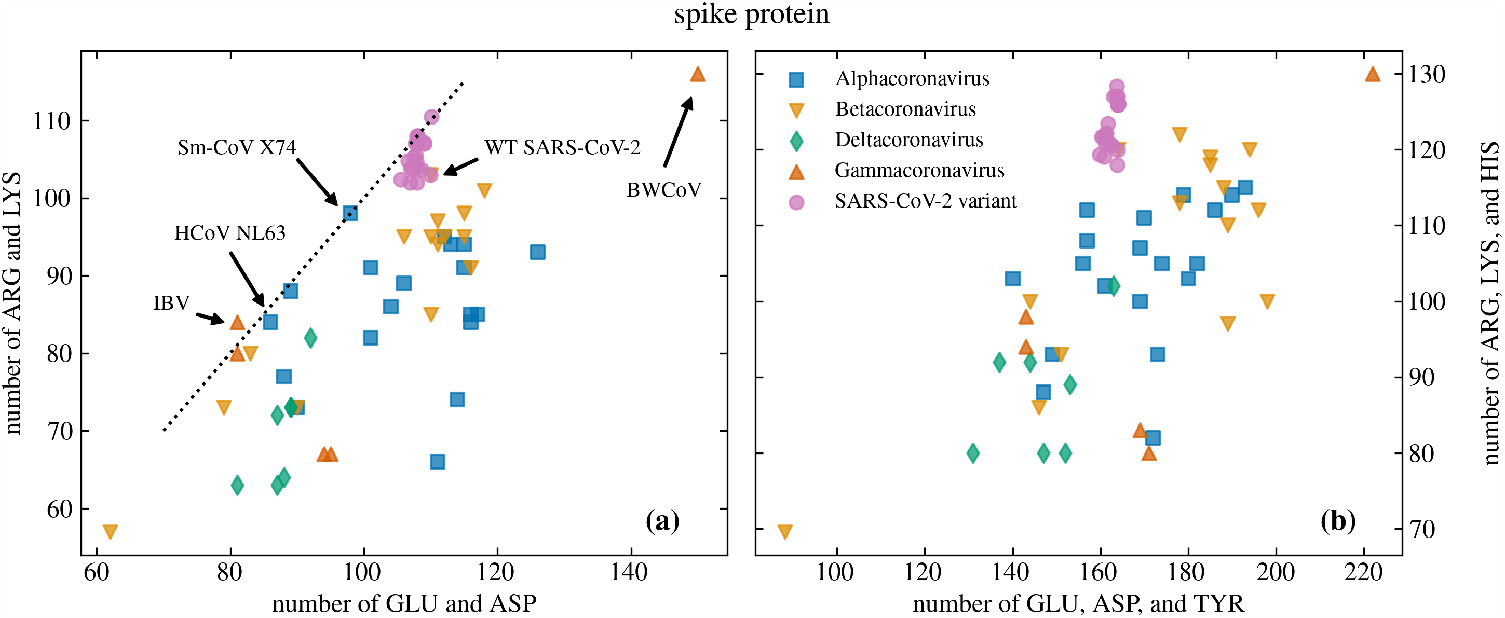
Comparison of the number of negatively and positively charged AAs on the S proteins of different coronaviruses. The comparison is shown for 46 different viruses from the Coronaviridae family and for 22 variants of SARS-CoV-2 (Table I). **(a)** The number of ARG and LYS compared to the number of ASP and GLU; **(b)** the number of ARG, LYS, and HIS compared to the number of ASP, GLU, and TYR. Dotted line in panel (a) shows where the number of negatively and positively charged AAs is equal. Arrows mark some of the coronaviruses mentioned in the main text.

## DISCUSSION AND CONCLUSIONS

By analysing the number of dissociable AAs on the S and N proteins of more than 1400 different SARS-CoV-2 lineages, we have shown that there is an overall tendency towards an increase in the number of positive AAs on the S protein with increasing lineage divergence, an effect which is not readily observed for the N protein (Fig. 3). While the number of dissociable AAs on a protein can only be considered a proxy for its total charge, which is in fact a result of many different local and global factors [22], our observations confirm that a more positively charged S protein of SARS-CoV-2 is evolutionarily favourable. This result is in line with previous studies [20, 21, 26] which have shown that the charge on the S protein has been increasing ever since the start of the pandemic. However, we have also shown that the overall number of positively charged AAs has seemingly reached a plateau with omicron and its subvariants (Figs. 4 and 1). Further observation of changes in the number of dissociable AAs in emerging variants as SARS-CoV-2 continues to evolve will show whether this plateau is only temporary or whether the positive charge on the S protein has reached its peak. On the other hand, our observation that the positive charge on the N protein, which plays a role in condensing the RNA genome [33–35], has changed only little with the evolution of SARS-CoV-2 might indicate that it is already optimized. We also note that while a similar analysis can be carried out on other structural proteins of SARS-CoV-2, such as the envelope and membrane proteins, these are significantly smaller than the S and N proteins and have far fewer dissociable AAs to start with. Consequently, any evolutionary changes in the number of dissociable AAs occur on a much smaller scale—on the order of a change in a single AA residue—and are thus difficult to compare.

Detailed inspection of the exact AA composition of the S protein in different SARS-CoV-2 lineages showed that the main contribution to positive charge comes from additional LYS residues, which reached the largest number with BA.1 and BA.3 omicron subvariants and the XD recombinant variant (Fig. 2a). Interestingly, the most divergent lineages, including the omicron subvariants BA.2.75 and BJ.1 as well as the XBB recombinant variant, show a significant increase also in the number of HIS residues, which is only fractionally charged at neutral pH. Comparison with the changes in the number of dissociable AAs on the N proteins of different SARS-CoV-2 lineages (Fig. 2b) shows that while there is a slight tendency for an increase in charge with increasing lineage divergence, this occurs to a far lesser extent. We argue that this is an additional confirmation that charge plays an important role for the function(s) of the S protein, and that the observed increase in the number of positively charged AAs on it is not a general effect that would occur in any viral protein as lineages continue to diverge.

Numerous studies have demonstrated how individual AA mutations that increase the local charge in the RBD of the S protein change its interaction with the ACE2 receptor (which is not the only receptor that binds the S protein of SARS-CoV-2 [36]). Even here, the question of positive charge is a complex issue: On the one hand, individual mutations which increase positive charge can reduce the binding affinity, as they are incompatible with particular LYS residues on the receptor [19]. On the other hand, the absence of charge-increasing Q493R mutation in the BA.4 and BA.5 omicron subvariants results in a significantly weaker binding affinity to ACE2 compared to omicron subvariants BA.1, BA.2, and BA.3 [17]. Charge can be an important factor also in other interactions, as it can disrupt the hydrogen bond network and thus influence the overall interaction [32], and it has the potential to diminish antibody binding [11]. Our results, together with previous studies [20, 21], clearly demonstrate that the evolutionary progress of SARS-CoV-2 lineages favours an *overall* increase in the positive charge of the S protein, making it stand out in this respect from among other known betacoronaviruses (Fig. 5). These changes likely have an effect which is greater than the contributions of individual AA mutations themselves: For instance, anionic, negatively charged lipids represent a dominant fraction of charged lipid species in biological membranes [37], and consequently their role in the interaction between proteins and membranes is of great biological interest [38]. More positively charged proteins would then interact more strongly with the membranes, consequently making the positive charge mutations more desirable. We hope that further studies can elucidate how the observed increase in the overall positive charge of the S protein benefits viral infectivity, transmissibility, and other traits.

## METHODS

### Data collection

#### SARS-CoV-2 variants

We obtained a list of SARS-CoV-2 Pango lineages from CoV-Lineages.org lineage report [39] on 24th November 2022. These lineages were used as an input to download virus genomic and protein data from NCBI Virus [40] using the provided command line tools; the data were downloaded between 30th November 2022 and 5th January 2023. We used the accompanying annotations to obtain the isolate collection dates and kept the earliest record with an available full date of collection (i.e., year, month, and day) as the timepoint of the lineage “emergence” for use in our analysis.

For each Pango lineage, we furthermore obtained the information on lineage divergence—the number of nucleotide changes (mutations) in *the entire genome* relative to the root of the phylogenetic tree, i.e., the start of the outbreak—from the global SARS-CoV-2 data available on Nextstrain [41]. We have selected only those entries with a genome coverage of > 99% and extracted their lineage divergence and the number of mutations. Since individual entries within a Pango lineage can still exhibit small differences in their lineage divergence from the WT genome, we averaged over them to obtain the average lineage divergence for each Pango lineage. To allow for an easier interpretation of our results, we list in Table I a comparison of the average lineage divergence of selected VOCs and VOIs used in our analysis.

As the last selection step, we retained only those Pango lineages whose downloaded protein fasta file was not empty. We used this protein data to obtain the number of dissociable AAs on the S and N proteins. The numbers of dissociable AAs were then averaged over all available protein data for a given Pango lineage. The final number of analyzed SARS-CoV-2 Pango lineages for which the entirety of the data described above was attainable is *N* = 1421.

#### Coronaviridae

As a point of comparison, we also examined the number of dissociable AAs on the S and N proteins of known coronaviruses. We used the coronaviruses listed in the most recent Virus Metadata Resource (2nd December 2022) issued by the International Committee on Taxonomy of Viruses (ICTV) [42], and limited ourselves to the genomes of those viruses with available REFSEQ accession numbers. We used these to download the representative genome and protein fasta files from NCBI Virus [40]. Due to the large amount of variation in the annotations across the different coronavirus datasets, we limited ourselves solely to the REFSEQ genomes and neglected any information on different strains and variants. This data was downloaded on 28th January 2023. We then followed the same procedure as with SARS-CoV-2 variants to obtain the number of dissociable AAs on the S and N proteins of different coronaviruses.

Our final dataset of coronaviruses includes 46 different species, of this one repetition of WT SARS-CoV-2. The dataset also includes SARS-CoV, MERS, and all other four known human coronaviruses. In general, the dataset comprises 19 species from the genus Alphacoronavirus, 15 species from the genus Betacoronavirus, 5 species from the genus Gammacoronavirus, and 7 species from the genus Deltacoronavirus.

#### Dissociable AAs

We analyzed the S and N proteins of SARS-CoV-2 variants and coronaviruses to obtain the (average) numbers of six dissociable AAs: glutamic (GLU) and aspartic acids (ASP), tyrosine (TYR), arginine (ARG), lysine (LYS), and histidine (HIS). We used Biopython to parse the protein fasta files and count the number of dissociable AAs on the S and N proteins. Three of the six AAs can carry positive charge (ARG, LYS, and HIS), while the other three can carry negative charge (ASP, GLU, and TYR) [43]. HIS typically carries a relatively small fractional charge at physiological pH, while TYR starts to acquire charge only at very basic pH; the importance of the latter in charge-mediated interactions has been demonstrated in recent studies [26]. In our analysis we did not consider cysteine (CYS), which has a thiol with a functional end group that is a very weak acid and is usually not considered to be an acid at all [44, 45].

## DATA AVAILABILITY

All the data presented in this work and a detailed description of data collection are openly available in OSF at https://osf.io/78b3f/, reference number 78B3F.

## ACKNOWLEDGMENTS

A.B. acknowledges support by Slovenian Research Agency (ARRS) under contract no. P1-0055. R.P. acknowledges funding from the Key Project under contract no. 12034019 of the National Natural Science Foundation of China.

## Notes

https://osf.io/78b3f/

### Competing Interest Statement

The authors have declared no competing interest.

### Summary of Updates

New figure added (Figure 1); revised parts of the MS to stress the importance of general, non-specific electrostatic interactions in the biological environment and especially in the context of SARS-CoV-2; minor clarifying changes in the Methods.

